# Look before you jump: jumping spiders discriminate different ants by visual cues

**DOI:** 10.1101/349696

**Authors:** Sajesh Vijayan, Chethana Casiker, Divya Uma

## Abstract

Ants, being ubiquitous, aggressive, and top predators, play a predominant role in terrestrial ecosystems. Jumping spiders are another prominent invertebrate predator that are present in similar habitats as that of ants. Most jumping spiders are thought to avoid ants, yet little is known if they discriminate among them. In this study we examined the response of jumping spider genus *Plexippus* towards three different ant species (*Oecophylla smaragdina*, the weaver ants; *Camponotus sericeus* the golden-back carpenter ants, and *Leptogenys processionalis*, the procession ants). In a behavioral assay that excluded tactile and chemical cues, we tested if spiders distinguish the three ants by visual cues alone. We recorded and analysed behaviors such as ‘look’, ‘approach’, ‘stalk’, ‘attack’, and ‘avoidance’ by spiders towards ants. Our results show that the three ants differ in their color, movement and aggressive behavior. Spiders gave ‘short looks’ to live ants, suggesting movement is important in detecting ants. Furthermore, spiders gave significantly more ‘long looks’ to procession and golden-back ants compared to weaver ants. Spiders approached, stalked and attacked procession ants more compared to weaver ants. Numerous jumping spiders and ants overlap in their habitat, and it is advantageous to selectively avoid some ants over others. Our results suggests that jumping spiders can indeed distinguish ants that co-occur in their habitat by visual cues alone, however, the precise nature of visual cues warrants further studies.

## Introduction

Ants play a prominent role in ecological communities. They are widely distributed, and top invertebrate predators in tropical ecosystems (Hölldobler, & Wilson, 1990; Roslin *et al.*, 2017). They compete for resources (including prey or nesting areas) with other arthropods, leading to non-consumptive effect on arthropod communities (Halaj *et al.*, 1997; Ibarra-Isassi, & Oliveira, 2018). Ants also serve as partners in mutualistic interactions, and their colonies sometimes serve as exploitable resources. Because of their ubiquitous presence, aggressive nature and communal defense, ants also serve as suitable models for numerous arthropod mimics (McIver, & Stonedahl, 1993; Cushing, 1998, 2012)

Since many invertebrate and vertebrate species have overlapping habitats with ants, and many even engage in various kinds of interactions with them, it may be advantageous to identify and distinguish among different ants. For example, coastal horned lizards selectively consume some ants over others and their preferences seem to be influenced by ant size (Suarez *et al.*, 2000). Butterflies avoid egg laying on plants frequented by predaceous ants, and this is influenced by ant size and shape (Sendoya *et al.*, 2009). Praying mantids avoid ant mimicking spiders because of their close resemblance to ants (Ramesh *et al.*, 2016).

Jumping spiders are another top arthropod predator in the terrestrial ecosystem, often found in similar habitat as that of ants. Earlier research suggests that many jumping spiders have an innate avoidance of ants or subsequently learn to avoid them (Edwards, & Jackson, 1994; Nelson *et al.*, 2006). However, it is not clear if jumping spiders can distinguish among different kinds of ants. Why might such a discrimination be important? Ants and jumping spiders overlap in their habitats, and are often intraguild predators (Okuyama, 2002; Sanders, & Platner, 2006; Sanders *et al.*, 2008). Furthermore, some ants may be more aggressive than others: For example, the weaver ant (*Oecophylla smaragdina*) is territorial and highly aggressive towards intruders. In such cases, it makes sense for spiders to employ a ‘safety first’ approach, and stay away from these ants. However, other ants may not be as aggressive, and jumping spiders could afford to forage closer to ant colonies, and sometimes even prey on insects tended by ants (Del-Claro, & Oliveira, 2000). In such cases, it may be beneficial for spiders to discriminate one ant from another. Jumping spiders may use various subcomponents of the visual signal such as movement, color and/or shape to distinguish between ants. Yet, the proximate visual cues these spiders use to discriminate different ants are not known. Multimodal cues used by these spiders in the visual, chemical or vibrational domain will have important implication for ecology and evolution of ant-spider interactions (Clark *et al.*, 2000; Allan, & Elgar, 2001; Nelson, & Jackson, 2006a), &, &.

The vision of jumping spiders is unique in the animal world, as they have achieved a high degree of spatial resolution due to their ‘principal eyes’ (Harland *et al.*, 2012), as well as a wide field of view due to ‘secondary eyes’ (Land, 1972; Forster, 1985; Jackson, & Pollard, 1996). Jumping spiders use shape, movement, and color to recognize a potential prey: for example, *Yllenus arenarius* rely on the direction of movement, and head-indicating features such as eye spots to strike a potential prey (Bartos, & Minias, 2016). *Evarcha culicivora*, which preys on female *Anopheles* mosquitoes appear to make discriminations based on relative orientation of body parts (Dolev, & Nelson, 2014). While movement is more essential than shape for predatory response in *Phidippus audax* (Bednarski *et al.*, 2012) the ant-eating jumping spider *Habrocestum pulex* seems to distinguish its prey even in the absence of motion cues (Li *et al.*, 1996). Both naïve and field-caught *Habronattus pyrrithrix* avoided red colored crickets (Taylor *et al.*, 2014), but they could also be trained to prefer them (Taylor *et al.*, 2016), demonstrating flexibility in color learning during foraging. Although these studies indicate the factors that influence prey recognition, cues that salticids use to identify a potential enemy/threat such as an ant have not been explicitly examined. Visual predators such as praying mantids avoided red colored ant mimicking spiders more than black colored ant mimics (Ramesh *et al.*, 2016). But, it is not known if salticids generalize all ants as dangerous or if they can distinguish between different species of ants.

The objective of the present study is to examine if jumping spiders can distinguish between three species of ants that co-occur in their habitat. We choose three ant species of comparable sizes: weaver ants (*Oecophylla smaragdina* Fabricius 1775), carpenter ants (*Camponotus sericeus* Fabricius 1798) and procession ants (*Leptogenys processionalis* Jerdon 1851) that differ in color, movement and aggression (Table 1). We first establish aggressive behavior, and movement of the three ants were indeed different. We then examined if jumping spiders differentiate these ants using vision alone by performing behavioral assays where tactile and chemical cues were controlled. Finally, by controlling for ant movement, we discuss the importance of color and movement cues influencing spider responses.

**Table 1:** Ants differed in color, aggressive behavior and movement.

| Ant | Color | Aggression | Movement |
| --- | --- | --- | --- |
| *Weaver ant* | Red | High | Continuous |
| *Procession ant* | Black | High | Continuous |
| *Carpenter ant* | Black-gold | Low | Sporadic |

## Methods

### Study species

Subadult and adult jumping spiders were collected from in and around Bengaluru, Karnataka from Oct 2016 to July 2017. After capture, the spiders were individually housed in plastic boxes with holes for ventilation, and a moistened piece of cotton for hydration. Spiders were captured two days prior to the behavioral assays. For the assays spiders in the genus *Plexippus* (*Plexippus petersi* Karsch 1878 and *Plexippus paykulli* Audouin 1826) were used, as they were predominantly found in the same habitat as that of ants. Foragers of all three species of ants and houseflies (used as control) were collected on the day of the experiment. Ant and representative spider pictures are shown in Fig. 1. All test animals were maintained and released back into their natural habitat in compliance with the laws of the country.

**Figure 1:**
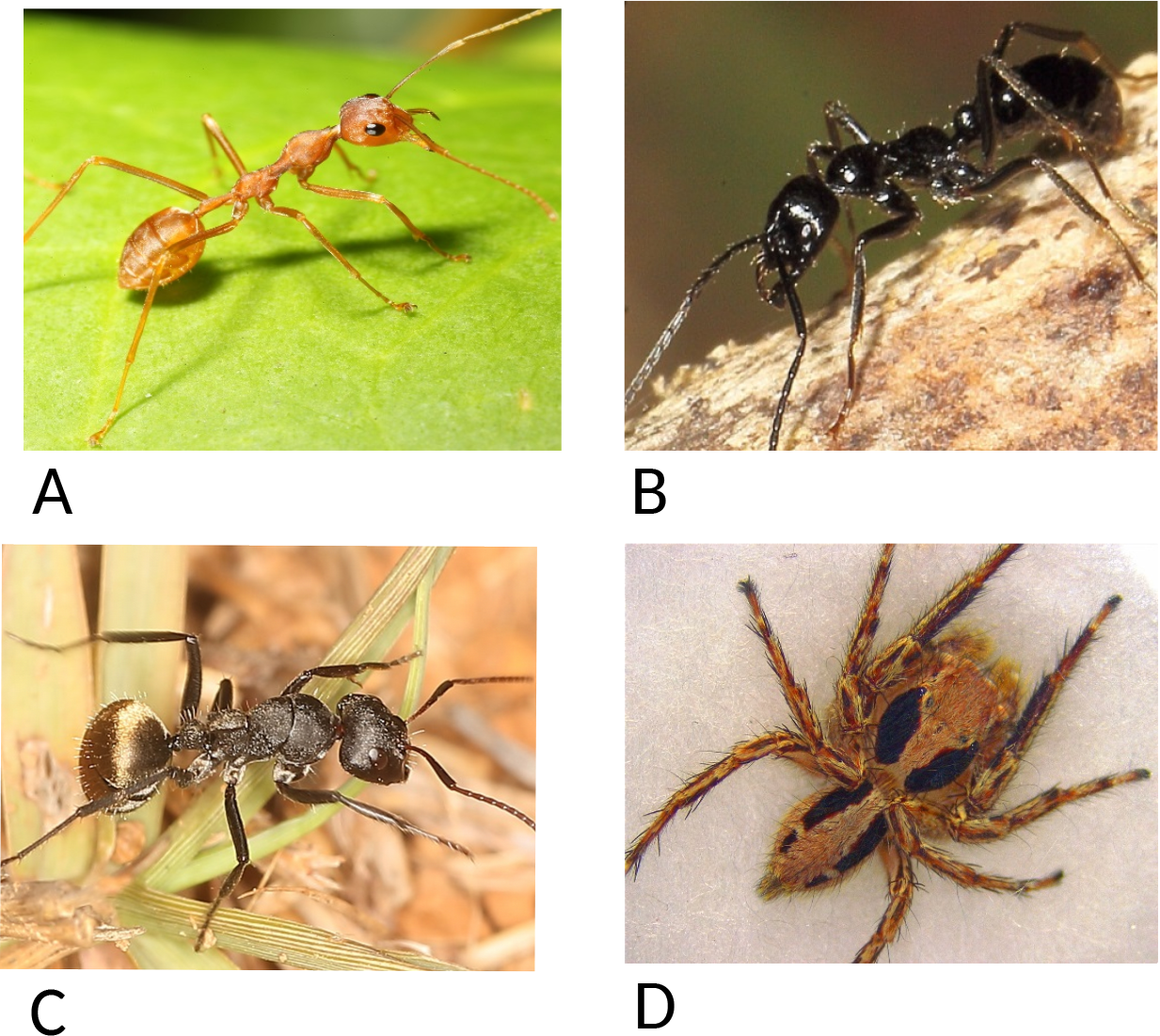
Three ant species and the jumping spider used in the study: A. Aggressive fast-moving weaver ants B. Aggressive, fast-moving procession ants C. Docile, staggered-moving carpenter ants D. *Plexippus petersi*, jumping spider (Photographs not according to scale)

### Ant aggression towards spiders

Ant aggression was calculated by restricting an ant (weaver, carpenter, or procession ants n=16 for each ants) and a spider in a 5 cm**^3^** arena and recording the latency of the ant to bite the spider. Ants which took more time to bite (increased latency) were considered to be less aggressive.

### Response of spider towards ants

We examined the response of spiders towards ants, where trials were carried out in an 8cm diameter Petri dish. The dish was modified to include an inner transparent ring made with a strip of transparent sheet into which the ant was released. The transparent ring neither allowed ants nor chemical cues to get into the outer compartment, thus allowing only visual cues to be seen from either side of the ring. An individual ant was introduced first, followed by a spider into the outer portion, upon which the trial commenced, and was video recorded using a Sony handycam (HDR-PJ600VE). At the end of 5 minutes, the trial was stopped and the test subjects were released back into the storage boxes. Trials were carried out from 9 am to 4 pm, well illuminated with a 4.5watt LED lamp. After each trial, the image of the spider was recorded using a Leica EZ4E microscope connected to a host computer, later used for identification. Each spider was presented with a weaver ant, carpenter ant or a procession ant (n = 29, n = 28 and n = 30 trials respectively). Using the same experimental setup, a different set of trials were carried out pairing a spider with a housefly to make sure spiders exhibited a typical hunting behavior (n = 25). This was to ensure that the set up did not hamper vision-mediated behavior of the spiders. Spiders or ants were used only once for all the trials.

Spider responses towards ants/flies were coded as follows: a) Short look: spider orients its cephalothorax towards the stimulus for a short amount of time (0-7 seconds) and then leaves b) Long look: spider orients its cephalothorax towards the stimulus for longer than 7 seconds and then leaves c) Approach: spider moving towards the stimulus, d) Stalk: spider moves slowly towards the stimulus, while visually oriented towards it e) Attack: spider pouncing on the wall of the outer ring at the stimulus f) Avoid: spider moving away from the stimulus. Short and long looks were used by Huang *et. al.* (2011) to classify jumping spider behavior towards different prey. Short looks in our study, on average, lasted for 1.5 secs (median 2.29 secs, range 0 to 7 secs) across all trials, and followed a beta distribution (Suppl Fig S1). Anything more than 7 seconds was considered long look, and it ranged from 7 to 112 seconds (average 20.3 secs, median 14.1seconds). The proportion of time spent and the frequency of these behaviors in 5 minutes was measured.

### Difference in ants’ movements

Ant movement was calculated as the amount of time the ants spent moving inside the inner transparent ring. The movement scores were obtained from the same trials where spider responses (see above) were scored. The presence of spiders outside the ring did not influence ant movement.

### Responses of spiders in the absence of ant movement

To control for the movement of ants, in a separate experiment, we created ‘lures’ of ants where freshly killed ants were positioned in a life-like manner inside the transparent barrier. Except movement cues, color and textural cues were intact. Spider responses (short and long looks) towards weaver, procession and carpenter ant lures were compared.

### Statistical analyses

We compared the aggression levels of three ants towards spiders, and movement of ants inside the ring by Dunn’s-test for multiple comparisons of independent samples with Bonferroni correction (Pohlert, 2014). We compared spider responses towards 1) weaver ant and carpenter ant 2) weaver ant and procession ant and, 3) carpenter ant and procession ant. The frequency of spider responses (approach, attack and avoid) and the proportion of time spent stalking and looking towards different species of ants were compared using Kruskal-Wallis test followed by post-hoc analysis with Dunn’s-test. Behavioral responses of the jumping spiders towards live ants and ant-lures; and house flies and ants were compared using Mann-Whitney U test. All statistical tests were carried out on R version 3.3.2 (R Core Team, 2018).

## Results

### Ants differ in their aggression towards spiders

Our results show that weaver ants, carpenter ants and procession ants differ in their aggression towards jumping spiders. Weaver and procession ants were very aggressive, and bit spiders in less than a minute after they were introduced as indicated by their average latency time to bite the spiders (Table 2). However, carpenter ants were not aggressive: 12 out of 16 trials did not elicit any aggressive response from the ants. Carpenter ants are observed to feed on extra floral nectaries and tend plant hoppers in the wild, and are known to be less aggressive (Mody, & Linsenmair, 2003).

**Table 2:** Aggression of ants, measured as latency to bite a spider. The more aggressive ants take less time to bite a spider. Both procession ants and weaver ants are aggressive compared to carpenter ants. Dunn’s test for ant aggression: Weaver ant vs Carpenter ant p = 0.007, Weaver ant vs Procession ant p = 1 and Procession ant vs Carpenter ant p = 0.001

| Ant | Latency to attack (sec) |
| --- | --- |
| *Weaver ant* (n = 16) | $63.32 \pm 68.67$ |
| *Procession ant* (n = 16) | $59.8 \pm 77.6$ |
| *Carpenter ant* (n = 16) | $156.9 \pm 52.1$ |

### Ants differ in their movement

Weaver ants and procession ants moved significantly more than carpenter ants when placed inside the ring (Table 3). While weaver and procession ants moved continuously, carpenter ants moved in a staggered manner, where they would stop and move multiple times. This behavior is similar to what is observed in the wild (pers. obs.).

**Table 3:** Percentage of time ants spent moving in experimental arena: Both weaver ants and procession ants move continuously, whereas carpenter ants moved sporadically. Dunn’s test for percentage of time spent moving by the ants: Weaver ant vs Carpenter ant p < 0.001, Weaver ant vs Procession ant p = 1 and Procession ant vs Carpenter ant p < 0.001

| Ant | Percentage of time spent moving |
| --- | --- |
| *Weaver ant* (n = 30) | 93.48 ± 9.69 |
| *Procession ant* (n = 29) | 86.41 ± 22.79 |
| *Carpenter ant* (n = 26) | 47.69 ± 38.54 |

### Spiders differentially respond to three ants

When controlled for aggression (by putting the ants inside a ring), the jumping spiders differentiated the ants by looks. Spiders gave quick short looks (median 2.29 sec) to all the three ants (Fig 2. Kruskal-Wallis *χ*^2^ = 1.337, df = 2, p-value = 0.5). On the other hand, spiders gave significantly longer looks (median 14.1 sec) to golden-backed carpenter and procession ants compared to weaver ants (Fig 3. Kruskal-Wallis *χ*^2^= 20.029, df= 2, p < 0.001. p values for post hoc Dunn’s test: carpenter ant vs weaver ant = 0.001, carpenter ant vs procession ant = 1 and procession ant vs weaver ant < 0.001). Furthermore, spiders approached, stalked and even attacked procession ants more than weaver ants. But these behaviors were not significantly different between carpenter ants and procession ants or carpenter ants and weaver ants (Table 4). Since ants were enclosed inside a barrier, they did not impose any threat towards spiders, and hence the spider sometimes approached and stalked these ants. When barrier was removed, the same spiders avoided these ants (unpublished results).

**Figure 2:**
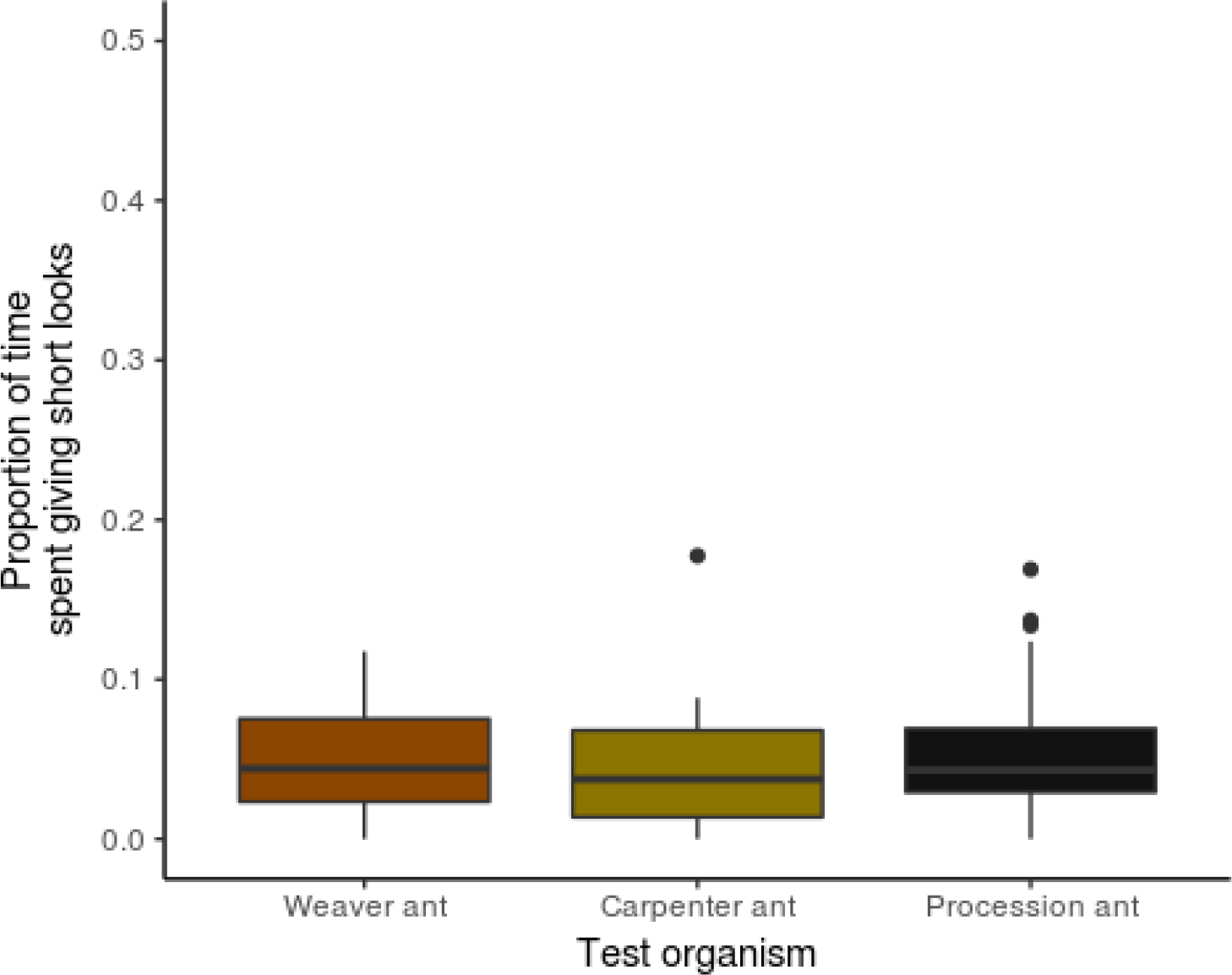
Proportion of time spider spent giving short looks: Proportion of time spent by spiders giving short looks to ants did not vary across the different ants. Kruskal-Wallis *χ*^2^ = 1.3377, df = 2, p = 0.5.

**Figure 3:**
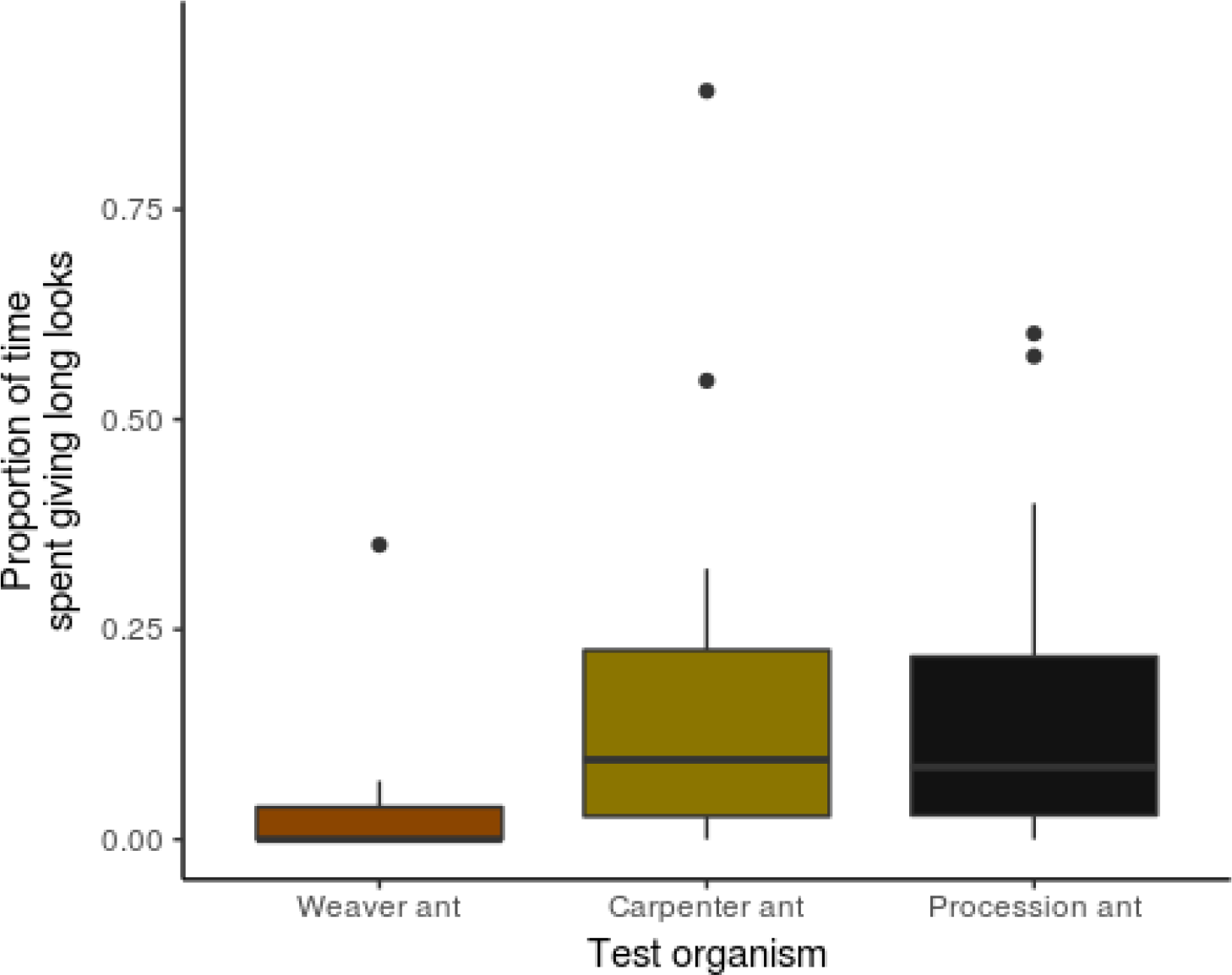
Proportion of time spider spent giving long looks: More time was spent giving long looks to procession ant and carpenter ants compared to the weaver ant. Kruskal-Wallis *χ*^2^ = 20.029, df = 2, p < 0.001. Post hoc Dunn’s-test: Weaver ant vs Carpenter ant p = 0.001, Weaver ant vs Procession ant p < 0.001, Procession ant vs Carpenter ant p = 1df = 2.

**Table 4:** Spider responses towards three ants: Spiders differed in their approach, stalk and attack towards the three ants (Kruskal-Wallis test: Approach: *χ*^2^: 7.45, P=0.02; Stalk: *χ*^2^: 10.13, P=0.006; Attacks: *χ*^2^: 6.35, P= 0.04; Avoid *χ*^2^: 0.51, P=0.7). Dunn Post hoc P values are presented below.

|  | *Weaver ant* vs<br>*Carpenter ant* | *Weaver ant* vs<br>*Procession ant* | *Procession ant* vs<br>*Carpenter ant* |
| --- | --- | --- | --- |
| Behavior |  |  |  |
| Approach | 0.29 | **0.02** | 0.20 |
| Stalk | 0.185 | **0.005** | 0.131 |
| Attack | 0.47 | **0.043** | 0.167 |
| Avoid | 1.000 | 1.000 | 1.000 |

Not surprisingly, jumping spiders, readily differentiated ants from the flies, corroborating that spiders can indeed differentiate prey from non-prey within our setup. When placed with a fly, all spiders exhibited a typical hunting behavior where they stalked (Mann-Whitney *U* = 665.5, p < 0.001) and attacked (Mann-Whitney *U* = 539.5, p < 0.001) the fly significantly more than they attacked any of the ants.

### Spiders responses in the absence of ant movement

In the absence of ant movement, spiders still looked at the ants, but did not differentiate them based on looks, suggesting that movement plays a crucial role in differentiating the three ants. Dunn’s test for looks by spiders towards lures gave non-significant results across all comparisons.

## Discussion

Most of the earlier studies suggest that jumping spiders avoid ants, but few have examined the response of spiders towards different ants. Our findings indicate that jumping spiders can distinguish and respond differentially to different species of ants. Weaver ants, procession ants, and carpenter ants differed in their color, aggression and movement, and spiders visually discriminated them. In this study we show that both movement and color of the ants influence the response of spiders.

Spiders when placed inside the Petri dish stayed predominantly at the periphery, and responded towards ants around 10-20 % of the time during which they differentiated between the ants. Our results suggest that movement of the ants is crucial for spiders to detect them. An earlier study had shown that jumping spiders innately avoided motionless dead lures of ants by visual cues alone (Nelson, & Jackson, 2006). Researchers found that spiders, at the end of 10 hrs, chose to stay away from the lures of ants devoid of chemical, movement, and textural cues. Although our study is not directly comparable to theirs, in the absence of movement cues the spiders failed to differentiate the three ants. Jackson, & Tarsitano (1993) found that out of 11 species of salticids, only one ant-specialist spider (*Corythalia canosa*) attacked motionless ant lures. Movement of an object, be it an enemy, prey or a mate is one of the primary cues that elicit spiders to orient towards the signal (Forster, 1977; Jackson, & Pollard, 1996; Bednarski *et al.*, 2012; Bartos, & Minias, 2016).

Our study suggests that while movement of ants elicit detection, other sub-components of visual cues (color, brightness, texture and/or behavior) enables spiders to distinguish between them. Spiders barely gave long looks to weaver ants compared to golden-back and procession ants. Additionally, our preliminary studies (not shown here) with seven other species (*Hyllus sp.*, *Menemerus sp.*, and from five other unidentified species) suggest a similar pattern. We have not explicitly tested whether spiders in our study perceive the red, black or black-gold colors of these ants, but recent studies have shown that spiders can see red (Zurek *et al.*, 2015), and avoid red colored prey (Taylor *et al.*, 2016). Regardless of *Plexippus* sp can see red, black or black-gold, it can nevertheless discriminate the ants.

While spiders seem to detect ants using short looks, they discriminate ants by giving long looks. What may be the significance of such a distinction? First, by giving frequent short looks spiders constantly scan their surrounding environment, keeping the target in its field of view. All three ants were moving (either continuously or in a staggered manner) inside the transparent ring, thus eliciting repeated short looks from spiders. Jumping spiders are known to continuously scan novel objects by looking for features like ‘legs’ or ‘head spots’ (Land, 1999; Bartos, & Minias, 2016). Moreover, jumping spiders in Huang *et al.*’s (2011) study gave more number of short looks to ants and ant mimicking spiders than non-mimetic spiders. Second, quick decisions have to be made when an enemy or a potential prey is dangerous or fast moving. For example, bumblebees make speedy decisions to avoid potentially dangerous foraging patches (Chittka *et al.*, 2009). Spiders may give frequent short looks to all ants and make a decision early on if it is a potential threat.

In addition to movement, spiders could use color to distinguish ants. Spiders gave more number of long looks to black colored ants; black color could belong to a potential prey or a predator. Spiders also approached, stalked and even attacked the black colored procession ants significantly more than the red colored weaver ants. This suggests that red colored ants could pose a stronger threat than black colored ones. Spiders’ response toward black-gold colored carpenter ants fell in between their response towards red and black ants. A recent study has suggested that gold/yellow color is also aposematic to a wide range of invertebrate and vertebrate predators (Pekár *et al.*, 2017). Whether color could play a significant component in distinguishing ants remains to be tested.

If predators with high visual acuity can differentiate ants, do they also respond differently towards different ant-mimicking spiders? Interestingly, praying mantids approached spiders that mimic black colored carpenter ants significantly more than the spiders that mimic red colored weaver ants (Ramesh *et al.*, 2016). That walking like an ant is indeed protective has been shown in recent studies (Pekár, & Jarab, 2011; Nelson, & Card, 2016; Shamble *et al.*, 2017). Whether visual predators respond to the mimics in the same way, by giving longer looks to some and short looks to others needs to be further tested.

There are over 5000 species of jumping spiders and most of them share habitats with different kinds of ants, especially in the tropics. Clearly it is advantageous to differentiate among ants rather than to avoid all of them and their mimics. How a spider perceives some ants as potentially dangerous is governed by dynamic interaction of multiple sensory modalities and contexts. We have shown the importance of visual cues in distinguishing among ants.

## Acknowledgement

This research was funded by a DBT BT/Bio-CARe/04/9809/2013-14, and APU grant to DU. Funding for CC and SV came from the DBT grant. We would like to thank Aranya Bagchi who helped collect spiders, and Dr. Deepak Deshpande for taking ant pictures.

## Contribution of authors

CC, SV and DU were involved in project design, data collection and analysis. SV and DU wrote the paper.

## Supplementary information

**Figure S1:**
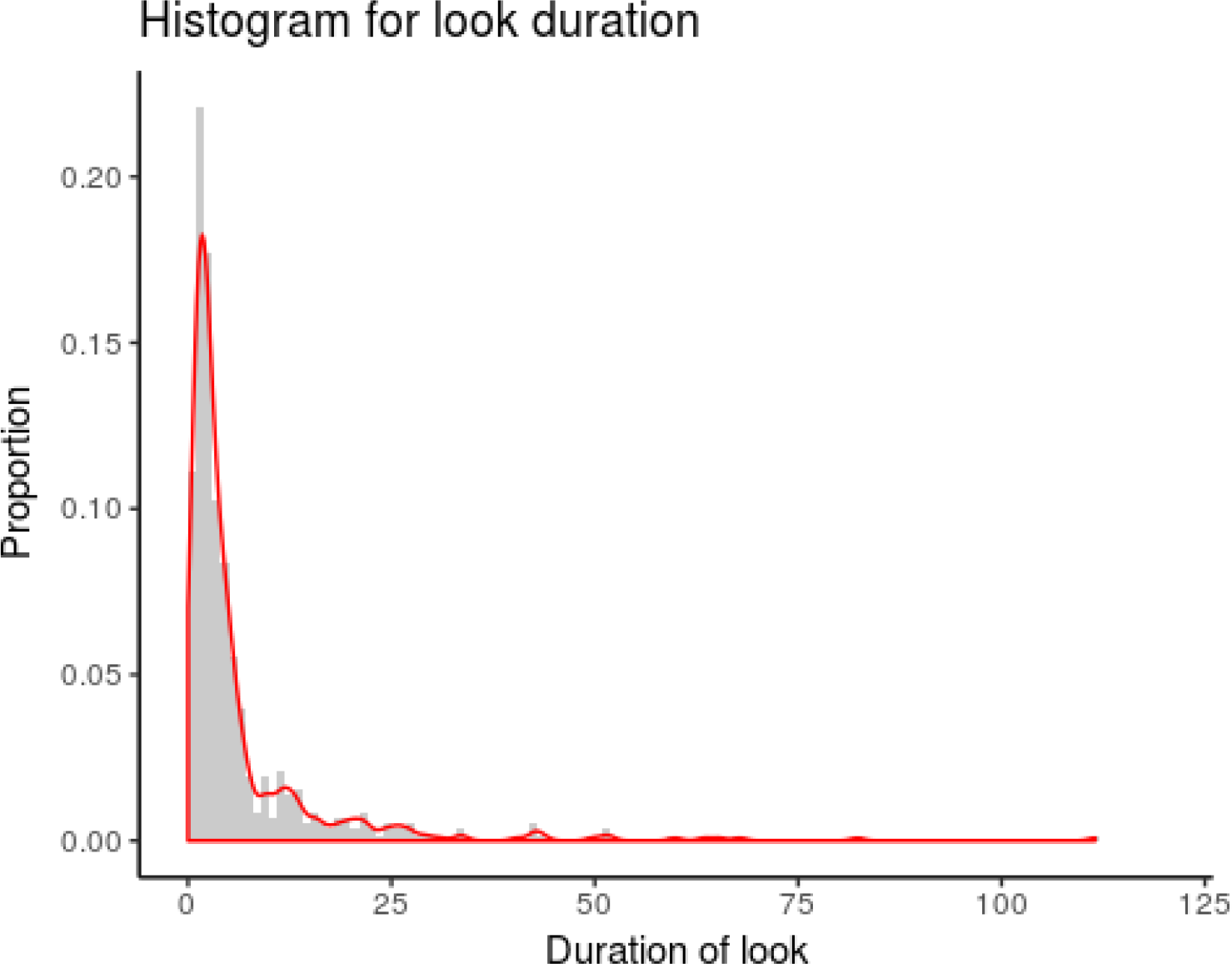
Histogram of duration of looks by spiders towards ants: The cut-off for differentiating between long and short looks has been established as 7 seconds. The normalized durations of short looks (duration less than 7 seconds) follow a beta distribution, with a median value of 0.02 (equivalent to 2.29 seconds).

**Table S1:**
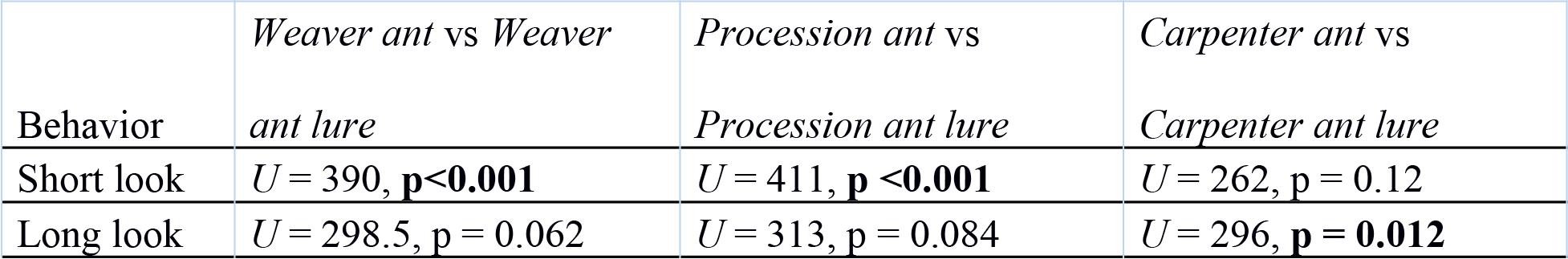
Spider responses towards live and dead ants: Spiders spent more time giving short looks to the live ants than their lure. Mann-Whitney U test scores comparing long and short looks of spiders towards ants and their lures are given below.

